# Loss of a morph is associated with asymmetric character release in a radiation of woodland salamanders

**DOI:** 10.1101/2025.02.12.637668

**Authors:** Brian P. Waldron, Maggie M. Hantak, Emily F. Watts, Josef C. Uyeda, Alan R. Lemmon, Emily Moriarty Lemmon, Robert P. Guralnick, David C. Blackburn, Shawn R. Kuchta

## Abstract

Color polymorphism, the occurrence of multiple discrete color morphs with co-adapted sets of traits within the same population, may provide the raw materials for rapid species formation. It has been hypothesized that fixation of a single morph can result in character release, whereby the monomorphic form evolves without the constraint of accommodating multiple adaptive peaks. However, the rates of evolution between populations fixed for different morphs likely depend on the specific adaptive zones occupied by each morph. We studied the evolution of dorsal color polymorphism (striped and unstriped morphs) in woodland salamanders (*Plethodon*), a North American radiation in which the polymorphism can be found in even the most distantly related species (∼44 Ma divergence). We estimated a phylogenomic tree of *Plethodon*, representing all extant taxa with multiple samples for most species. Morphometric data suggest that between-species variation exists predominantly along an axis of relative body elongation, likely corresponding to a terrestrial–fossorial continuum. Polymorphic species occupy an intermediate phenotypic space between the evolutionary optima of striped and unstriped species, although polymorphic species did not have elevated speciation rates. Faster rates of body shape evolution were observed in unstriped species, suggesting that body elongation, which is co-adapted with the unstriped morph, is constrained by the polymorphism. Striped species had slower rates of evolution than polymorphic species, despite lacking the genetic constraints often associated with polymorphism. Our results demonstrate that rates of phenotypic evolution and speciation following character release can be asymmetric and idiosyncratic depending on the alternative adaptations of each morph.

## Introduction

Phenotypic polymorphisms—discrete, genetically-based variants within a single interbreeding population, the rarest of which is too common to be maintained by recurrent mutation—can be shared or sorted across species in an evolutionary radiation (1–3). Morphs comprise unique combinations of co-adapted traits that are maintained by multivariate disruptive selection, effectively allowing one species to occupy multiple niches within a population (2, 4). Because polymorphisms provide divergent adaptive phenotypes for natural selection to act upon, it has been hypothesized that they can also promote rapid speciation following loss of one or more morphs (5, 6). This process has been identified by West-Eberhard as a type of character release (7, 8), as the loss of one morph can enable a population to evolve at faster rates towards a phenotypic optimum by eliminating or reducing the genomic constraints associated with accommodating multiple adaptive peaks within a single population. Thus, polymorphism may facilitate diversification, and studies have reported increased speciation rates in color polymorphic birds (9), lizards (5), and carnivores (10). However, rates and patterns of phenotypic evolution following character release can differ between populations or among species. Depending on each morph’s underlying ecological, genetic, and developmental background, rates of phenotypic evolution and species formation may also be asymmetric: that is, fixation of a morph may result in evolutionary stasis for some morphs, but rapid evolutionary change in others (11, 12).

For this study, we focused on the evolution of a conserved color polymorphism in woodland salamanders (family Plethodontidae; genus *Plethodon*), a classic species radiation in North America (13). Many species of *Plethodon* have a dorsal color polymorphism, with populations containing some individuals with a red dorsal stripe (“striped”), while others lack the stripe and are entirely black (“unstriped”; Fig. 1) (14, 15). The polymorphism is common across *Plethodon*, including western North American (subgenus *Hightonia*) and eastern North American (subgenus *Plethodon;* Fig. 1) clades (16). Moreover, most monomorphic clades have been hypothesized to descend from polymorphic ancestors that lost one of the two color morphs (17), and developmental data suggest that all species, including those completely unstriped, have a stripe during the embryonic or juvenile period (18–21), providing a potential mechanism for transitions among color morph states.

**Figure 1.**
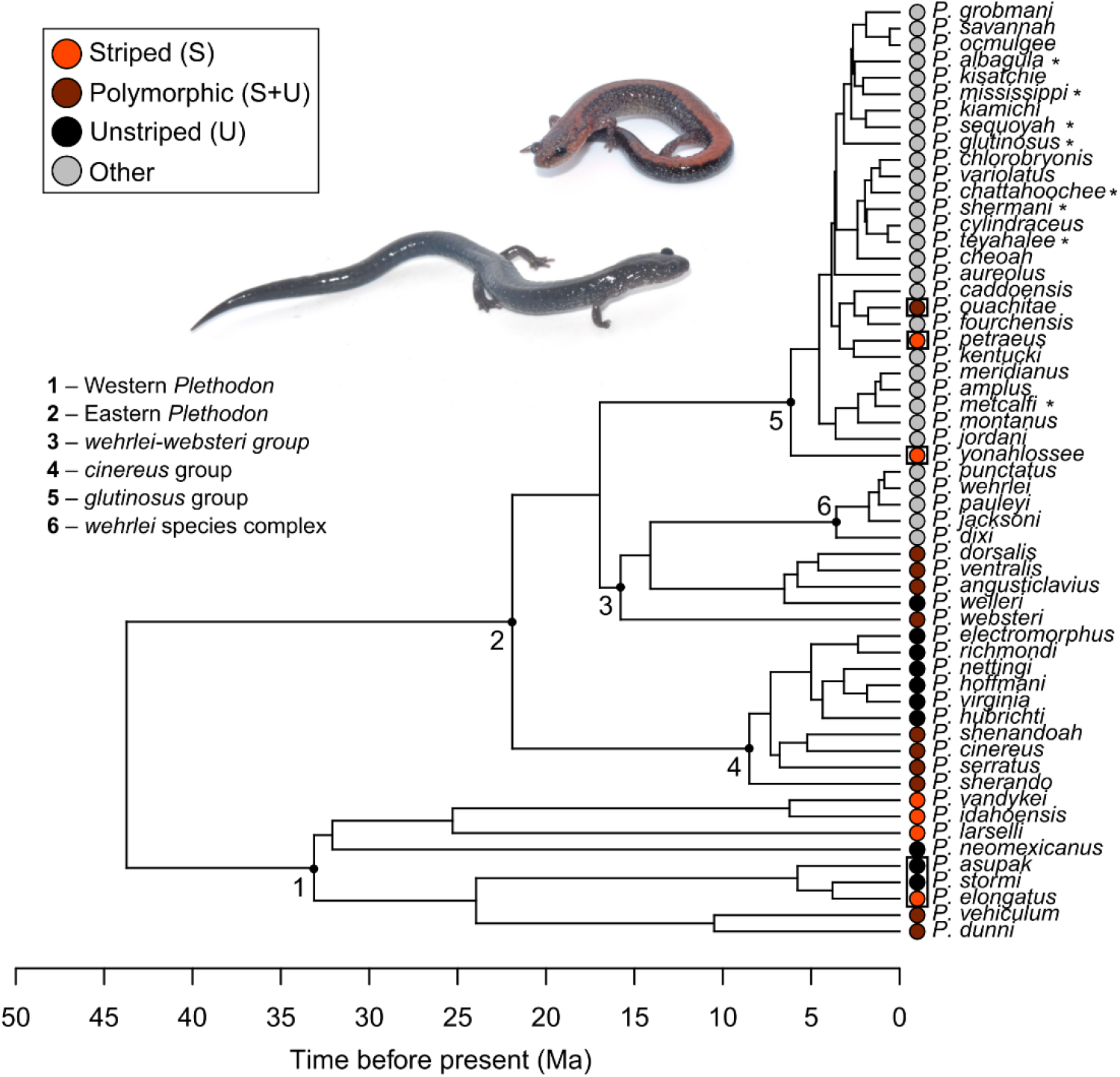
Timetree of *Plethodon*. Several clades recognized in the manuscript are numbered, and tip colors indicate whether the species is striped (S), unstriped (U), or polymorphic (S+U) for the dorsal color pattern. Most large-bodied species in the *glutinosus* group and *P. wehrlei* species complex were considered a separate color category from striped/unstriped (“other”). Boxes show species for which we tested alternative color classification schemes (see text). The species tree topology and divergence dates were estimated using wASTRAL and MCMCTree, respectively. Species with multiple phylogroups, or possibly paraphyletic or polyphyletic species (*), were pruned to a single representative sample. Photos show a striped morph of the polymorphic species, *P. cinereus* (top), and a fixed unstriped species, *P. hoffmani* (bottom).

In polymorphic species of *Plethodon* (particularly the well-studied Eastern Red-backed Salamander, *P. cinereus*), several lines of evidence—including field and laboratory experiments and macroecological studies—suggest that color morphs exhibit ecological divergence along several axes, including crypsis (22), diet (23, 24), climatic niche (17, 23, 25–28), and behavior (29–31). Morphologically, unstriped species tend to have more trunk vertebrae, which may correspond to body elongation and a more fossorial ecology (17). However, it is unclear how the color morphs differ in other key body shape characteristics (i.e. limb, head, and body allometry), whether polymorphism is a necessary “stepping stone” for transitions between striped and unstriped states, whether fixation of a morph is correlated with different rates or patterns of morphological evolution or species diversification, and if large-bodied species of *Plethodon* (i.e., the *glutinosus* group and *P. wehrlei* species complex; Fig. 1), which have different color patterns and body shape allometries compared to small-bodied species, evolve at different rates than the smaller striped/unstriped species.

To address these questions, an accurate characterization of the phylogenetic history of *Plethodon* is needed. Previous studies have employed datasets that included up to 12 loci (32), although several species, including some recently described (33–35), were not sampled (13, 32, 36). Furthermore, phylogenetic studies commonly include only a single sample for most species, while including multiple geographic representatives along with many more loci can provide more robust estimates of relationships and identify cases of non-monophyletic taxa (37, 38). In this study, we inferred a well-sampled phylogenomic tree of the North American radiation of *Plethodon* salamanders, including representatives of all extant species, to address hypotheses about morphic evolution. In particular, we use our phylogenomic tree to test whether speciation tempo and mode was correlated with the dorsal striped/unstriped color polymorphism. We hypothesized that: 1) monomorphic species or clades result from the loss of a morph present in a polymorphic ancestor; 2) monomorphic species have faster rates of phenotypic evolution than polymorphic species, due to release from genomic constraints associated with polymorphism; 3) monomorphic striped and unstriped species have different evolutionary optima consistent with alternative adaptive sets of traits; 4) polymorphic species have faster speciation rates than monomorphic species; and 5) large-bodied species that lack the dorsal stripe—those in the *glutinosus* group and *P. wehrlei* species complex—have lower rates of phenotypic evolution consistent with the interpretation of a “non-adaptive” radiation of morphology in these groups (13, 39).

## Results

### Phylogenomics of *Plethodon*

We recovered a largely well-resolved phylogeny using 282 anchored hybrid enrichment loci from all extant species of *Plethodon* (57 species; N = 194 ingroup samples), including multiple samples for all eastern *Plethodon* (Fig. 1; SI Appendix, Table S1, S2). Some species that included phylogeographic variation or were not monophyletic in our concatenated analysis (IQ-Tree; SI Appendix, Fig. S1) were inputted as multiple operational taxonomic units for species tree estimation (wASTRAL; SI Appendix, Fig. S2). Following time-calibration, each species was pruned to a single representative based on the current taxonomy for downstream comparative analyses (Fig. 1; details in SI Appendix). All phylogenies, including species trees with a single tip per species or with multiple units included for some species, are available as Supplemental Data.

### Color morph evolution and asymmetric character release

We divided species of *Plethodon* into color morph categories of striped (S), unstriped (U), or polymorphic (S+U). An additional category (“other”) was used for most or all species in the *glutinosus* group and the *P. wehrlei* species complex, which are larger-bodied species lacking the striped/unstriped phenotype. We created eight alternative classification schemes to account for uncertainty for some species (boxes in Fig 1; see SI Appendix, Table S3). Then, we compared 12 models of discrete character evolution (Mk models) that varied in the direction and symmetry of transitions among color morph states, including several models that explicitly considered polymorphism as an intermediate step between monomorphic states (SI Appendix, Fig. S3). The ancestor of eastern *Plethodon* was reconstructed as likely polymorphic (posterior probabilities ranged 0.58–1.0 across schemes), while the ancestors of all *Plethodon* and of western *Plethodon* were uncertain or sensitive to the classification scheme (Fig. 2; SI Appendix, Fig. S4). Based on AIC scores across models, most schemes (6 of 8) supported a directional polymorphism model, in which transitions occurred asymmetrically from striped to polymorphic, and from polymorphic to unstriped (SI Appendix, Table S4). However, the best-fitting models of discrete character evolution in these analyses were sensitive to the classifications. Other schemes supported a simpler equal-rates model between any state, and when directional models were favored, AIC values were typically similar (ΔAIC < 2.0) for other models (i.e. variants of equal rates, symmetrical rates, or “dead end” models in which all transitions to fixed striped or unstriped were asymmetrical from polymorphic; SI Appendix, Table S4). Even so, in eastern *Plethodon*, which had the greatest certainty in color classification, there were two well-supported cases of fixation for the unstriped morph from polymorphic ancestors, whereas no cases existed for fixation of the striped morph (Fig. 2; SI Appendix, Fig. S4).

**Figure 2.**
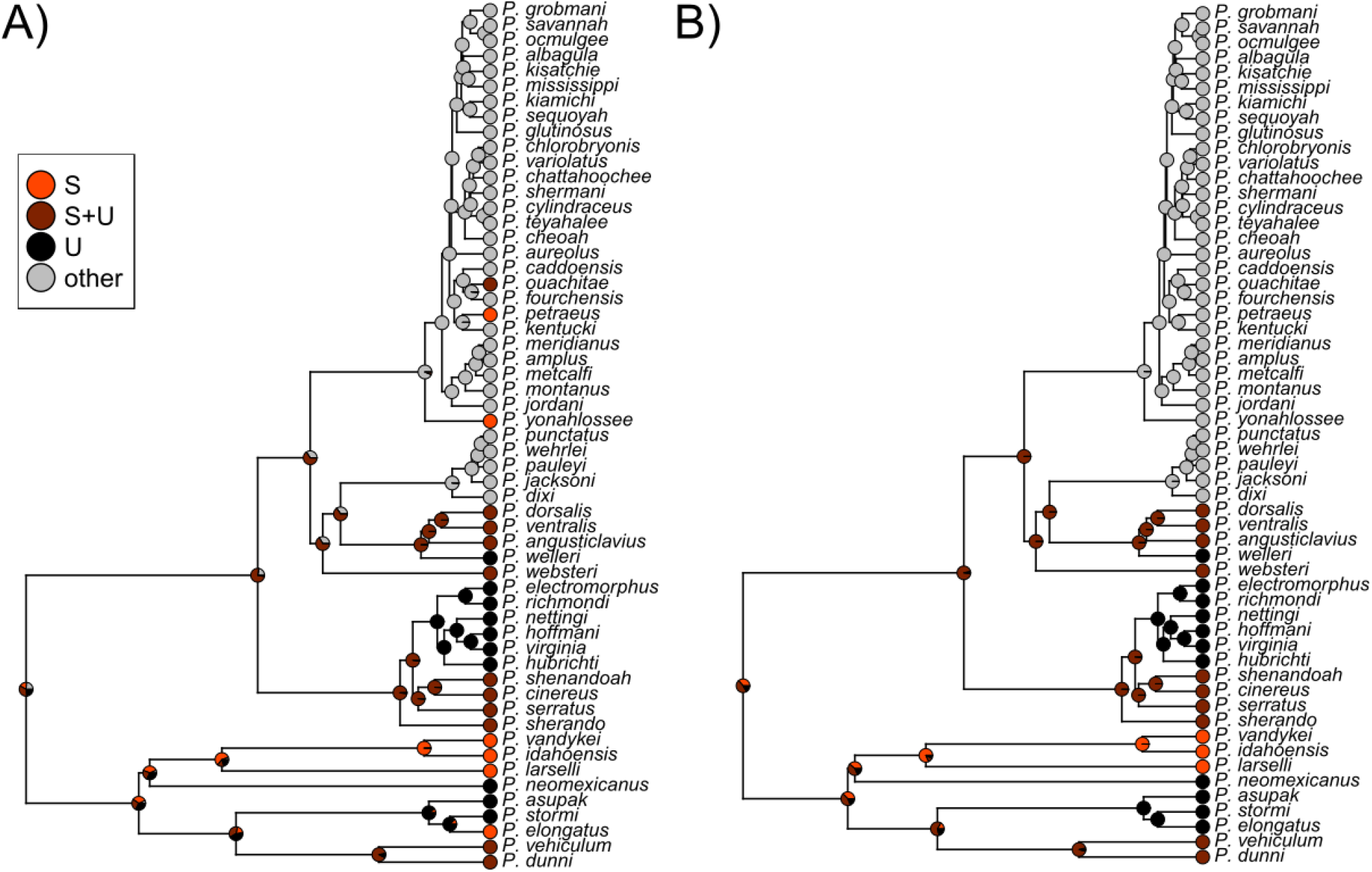
Ancestral character reconstructions of color morphs comparing two color classification schemes, classifying each species as striped (S), unstriped (U), polymorphic (S+U), or other. In (A), tip states match those shown in Fig. 1, whereas (B) shows an example of an alternative scheme in which all members of the *glutinosus* group were classified as “other,” and the western species *P. elongatus* was unstriped. Pies at internal nodes represent the results of 1,000 stochastic character maps. Most estimated ancestral states were consistent across schemes; uncertainty within and between schemes was greatest at the root and within western *Plethodon*. Reconstructions for all classification schemes are shown in the SI Appendix (Fig. S4).

All morphometric traits—raw costal groove count (CG) and body-size-corrected forelimb length (FLL), hindlimb length (HLL), head length (HL), snout-eye distance (SE), and body width (BW)—exhibited significant phylogenetic signal (*p* < 0.05), with lambda >0.90 for all traits except head length (lambda = 0.80; SI Appendix, Table S3, S5). Variation in costal groove count (i.e., trunk vertebrae) and continuous traits largely corresponded to relative elongation (phylogenetic PCA; Fig. 3a, SI Appendix, Fig. S5), in which elongated species had more trunk vertebrae and proportionally shorter limbs, heads, and snouts (pPC1; 58.3% of variance explained); the next highest axis of variation corresponded mostly to body width (pPC2; 18.1%; SI Appendix, Table S6).

**Figure 3.**
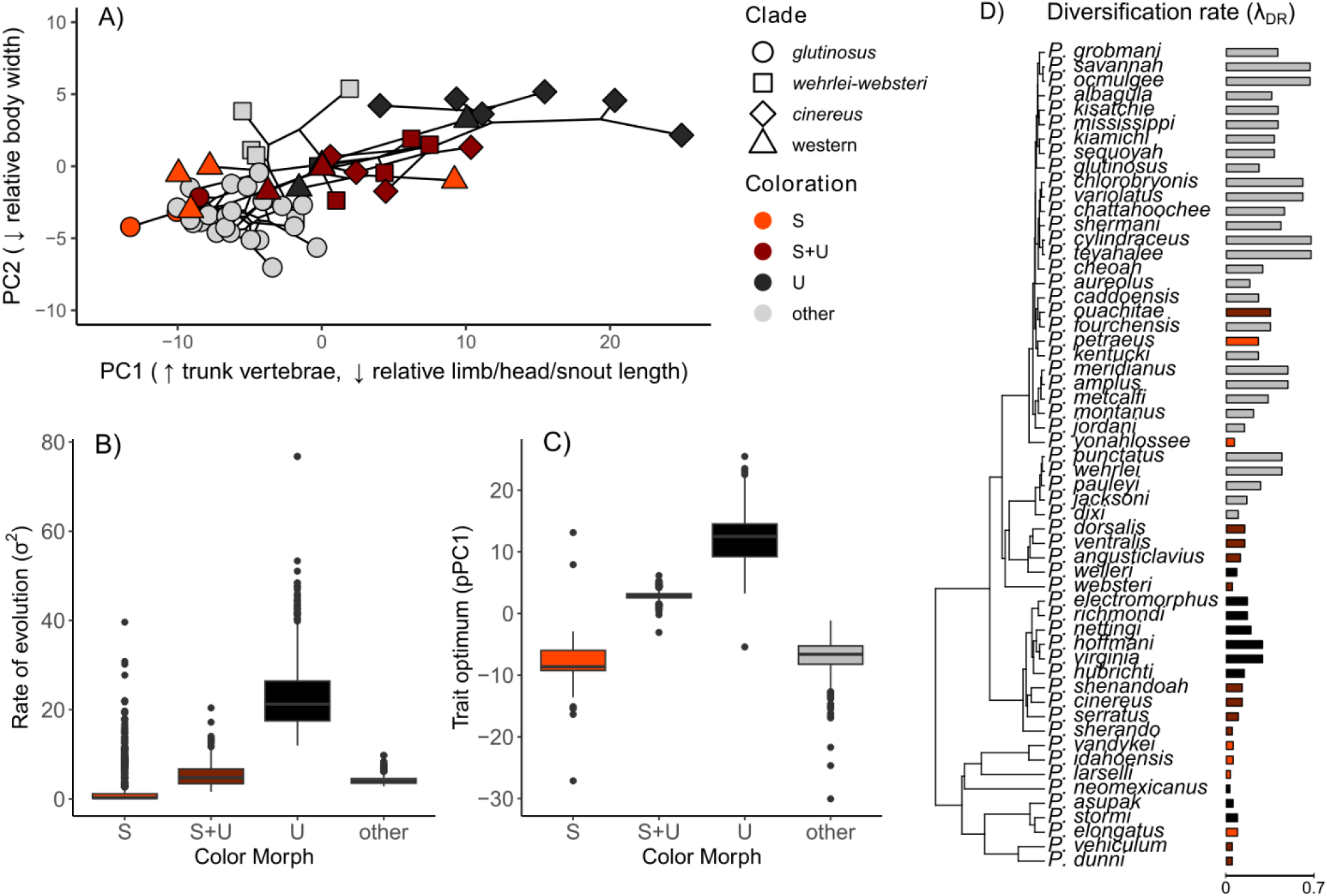
Unstriped species are more elongated and evolve more rapidly than polymorphic species. A) Phylogenetic PCA (pPCA) of morphometric characteristics across species. The first axis is consistent with increasing numbers of trunk vertebrae and decreasing body-size-corrected lengths of the limbs, head, and snout. The second axis represents decreasing size-corrected body width. Points are labeled by species groups and colored by dorsal color state (S = striped, S+U = polymorphic, U = unstriped). Represented species colorations match Fig. 1, and a version of the figure with species names labeled is available in the SI Appendix Fig. S5. B) Rates of evolution and C) trait optima estimated using OUwie for pPC1, an axis positively related to relative body elongation. Results across all coloration schemes and stochastic character maps are combined in boxplots for each color morph state. D) Diversification (i.e. speciation) rates tended to be higher in unstriped species, a pattern driven by an unstriped clade in the *cinereus* group. Species in the “other” category had the highest rates overall. Represented species morph classifications also match with panel (A) and Fig. 1.

Unstriped species displayed faster rates of phenotypic evolution and more elongated morphologies than polymorphic species (multi-rate and optima OU models in OUwie; Fig. 3b, c; Table 1; SI Appendix Fig. S6). Striped *Plethodon* typically had a less elongated morphology than polymorphic species—consistent with a less fossorial adaptive zone—yet their estimated rates of evolution were as slow or slower than polymorphic species (Fig. 3b, c; Table 1). Large-bodied (“other”) species, including the diverse *glutinosus* group, did not mirror patterns of evolution in unstriped species (despite most species lacking a red dorsal stripe); instead, their estimated rates of evolution were lower and most similar to polymorphic species, and they had a lower optimum for body elongation that was most similar to striped species (Fig. 3b, c; Table 1). Results were qualitatively similar across color morph classification schemes and whether pPC1 (Fig. 4) or univariate measurements of relative elongation were used (SI Appendix, Fig. S7).

**Table 1.** Rates of phenotypic evolution and lineage diversification rates across color morph states. Phenotypic optima (θ) and rates (σ^2^) were estimated using OUwie for the first axis of phylogenetic PCA, representing mean values and standard errors (SE) for 100 stochastic character maps repeated across eight color classification schemes. Lineage diversification rates, or tip rates, were estimated using the DR statistic (λ_DR_). Mean λ_DR_ is the mean value for species across all eight classification schemes, while the mean range provides the lowest and highest mean values across schemes. The min λ_DR_ and max λ_DR_ represent the minimum and maximum tip rates, respectively, observed across species within each color morph state for any scheme.

|  | Phenotypic evolution |  |  |  | Lineage diversification |  |  |  |
| --- | --- | --- | --- | --- | --- | --- | --- | --- |
| | Optimum ($\theta$) | SE | Rate ($\sigma^2$) | SE | Mean $\lambda_{DR}$ | Mean range $\lambda_{DR}$ | Min $\lambda_{DR}$ | Max $\lambda_{DR}$ |
| Striped | -7.7 | 0.077 | 2.3 | 0.164 | 0.073 | [0.048 – 0.093] | 0.033 | 0.258 |
| Polymorphic | 2.8 | 0.024 | 5.5 | 0.087 | 0.106 | [0.092 – 0.119] | 0.048 | 0.355 |
| Unstriped | 12.0 | 0.133 | 23.7 | 0.283 | 0.161 | [0.146 – 0.172] | 0.028 | 0.291 |
| Other | -6.6 | 0.079 | 4.2 | 0.029 | 0.391 | [0.383 – 0.399] | 0.064 | 0.682 |

### Lineage diversification among color morphs

Patterns of diversification rates (hereafter, speciation rates) among color morph states were similar to those of phenotypic evolution: unstriped species tended to have higher tip speciation rates (λ_DR_) than polymorphic or striped species, a pattern driven by diversification of unstriped species within the *cinereus* group (Fig. 3d; Table 1). Striped species had similar or lower rates than polymorphic species (Table 1), although we interpret these results cautiously given our tree size, particularly for striped species. Large-bodied (“other”) species, despite having relatively low rates of phenotypic evolution (Fig. 3b), had the fastest speciation rates, with an average rate over twice that of unstriped species (Fig. 3d; Table 1). Taken together, these results suggest that polymorphism is not clearly associated with faster speciation rates in *Plethodon*.

## Discussion

The theory of character release predicts that fixation of a morph from a polymorphic ancestor can result in faster rates of phenotypic evolution and a shift towards phenotypic optima (7). In *Plethodon*, we found strong evidence of these predictions for the unstriped morph and partial evidence for the striped morph. Species fixed for the unstriped morph experienced elevated rates of phenotypic evolution and an increase in body elongation. By contrast, species fixed for the striped morph had the lowest rates of phenotypic evolution, a result that was robust to uncertainty in color morph assignment in our analyses. Striped species did, however, occupy a distinct phenotypic optimum (i.e., shorter relative length) compared to both unstriped and polymorphic species, while the optimum of polymorphic species was intermediate. These results suggest that polymorphic species straddle distinct phenotypic optima between monomorphic states.

Character release in *Plethodon* occurs along an axis of elongation, where unstriped species tend to be relatively longer in body length with reduced relative head and limb size while striped species have relatively stouter bodies. These characteristics are commonly associated with a spectrum from fossoriality (more elongated) to terrestriality (less elongated) in a variety of vertebrate taxa (40–43). There are multiple possibilities for why the unstriped phenotype is associated with elongation and fossoriality. Experimental studies involving visual predators suggest that negative frequency-dependent selection could maintain the polymorphism within species (27, 49, but see 46), particularly if the morphs differ in their time spent above and below ground. The striped morph may be better camouflaged in a terrestrial environment of leaf litter or woody debris, whereas the dorsal stripe may lose its adaptive significance in fossorial environments (22), resulting in disruptive selection for dorsal coloration. It is also possible the unstriped condition is genetically correlated with other strongly selected traits involved in microhabitat use (4, 47), such as those related to thermal or hydric physiology (48, 49). Past work has suggested divergent climate niches for the morphs that could correspond to fossoriality/terrestriality (23, 25, 27), although the generality of these patterns remains unclear because most experimental work has focused on a single species, *P. cinereus* (17, 26).

The elongated form associated with fixed unstriped species experiences faster rates of phenotypic evolution than polymorphic or fixed striped species. This pattern is consistent with release from genomic constraints when a morph is lost (7, 8). The unstriped morph may also have greater phenotypic space to explore because elongation can be achieved by adding vertebrae (or, in rare cases in the genus *Pseudoeurycea*, by lengthening vertebrae; 41), whereas there is a sharper lower limit for the less elongated striped morph (50). For example, no salamander has fewer than 12 trunk vertebrae, while some species possess up to 61 (36).

Asymmetric evolutionary rates and optima between phenotypes also call into question the lability of fixation for color morphs, particularly in more extreme elongated forms. In our analyses, morph loss was biased towards fixation of the unstriped morph; for example, eastern *Plethodon* had at least two transitions from a polymorphic ancestor to the monomorphic unstriped state, yet no transitions from polymorphic to striped were inferred. This is notable considering that the striped morph is more common in all species that have both morphs, with species-level estimates ranging from 69.3–98.7% (17). Species of all major clades develop a stripe at some point during ontogeny (18–21), suggesting that while the stripe should be possible to maintain or re-evolve, fixation of the unstriped morph may be due to positive natural selection.

Color polymorphism is associated with higher speciation rates in several other taxa (5, 9, 10). If the fixation of a morph is followed by strong directional selection toward a phenotypic optimum, polymorphism can promote speciation. Morphs represent contrasting character sets produced by a single genome, allowing a population to occupy multiple niches (7). Morph fixation releases constraints on the genome related to the production of multiple phenotypes, allowing the species to reach a more fully optimized phenotypic state. For example, in the classic “rock-paper-scissors” model of alternative reproductive tactics in the Side-blotched lizard, *Uta stansburiana*, male morphs are associated with three throat colorations (orange, yellow, blue) that are correlated with distinct mating strategies (51). Loss of a morph in this system is linked with rapid evolutionary change in body size, which may promote the formation of incipient species if the evolution of body size or other heritable traits reinforces reproductive isolation between populations (8). However, polymorphism can also promote speciation more generally by expanding the species’ ecological niche breadth if, for example, wider niches lead to wider geographic distributions followed by range fragmentation (39, 52, 53). These processes may leave distinct signals in the phylogeny related to polymorphism and speciation rates. In the case of character release, polymorphism may be associated with faster speciation, but with a bias towards monomorphic daughter lineages, as the fixation of a morph has a causal relationship with phenotypic divergence and speciation (8, 10). Alternatively, in the case of expanded niche breadth, polymorphism should be associated with faster speciation in polymorphic lineages more generally (10).

Polymorphism in *Plethodon*, however, was not clearly associated with faster rates of speciation. Speciation rates tended to mirror patterns of phenotypic evolution, whereby unstriped species had the fastest rates, striped the slowest, and polymorphic species were either intermediate or similar to striped species (with some uncertainty due to the size of our phylogeny). The fact that unstriped species, specifically those in the *cinereus* group, are associated with a more fossorial morphology and have faster speciation rates contrasts with patterns among highly fossorial clades of amphibians, which are typically less species-rich (54, 55). However, salamanders that live on land—those that are not aquatic or arboreal—are rarely fully terrestrial or fully fossorial, but instead exist along a continuum of terrestriality to semi-fossoriality (55, 56). Our results suggest that species that are more fossorial can evolve more quickly in both morphology and species diversification, a result that should be tested in other clades with similar microhabitat use (41, 42).

The *glutinosus* group and *P. wehrlei* species complex—all large-bodied species that we treated as a separate morph category (“other”)—showed a dissimilar relationship between lineage diversification and rates of phenotypic evolution compared to striped/unstriped species. Despite exceptionally low rates of morphological evolution, the *glutinosus* group and *P. wehrlei* complex had the highest speciation rates. These groups likely evolved from polymorphic ancestors, and therefore the evolution of large-bodied clades may be consistent with character release and diversification following fixation of a morph, as observed in unstriped species. Phenotypic evolution, though, remained similarly slow to that of polymorphic species. This result is consistent with previous comparative analyses of lungless salamanders (family: Plethodontidae) showing that rates of morphological evolution tend to be decoupled from diversification rates (57). Our results also highlight a critical context-dependence of character release from the constraints of polymorphism: increased rates of morphological evolution or diversification are not inevitable consequences of fixation of a morph, but rather they can depend on the adaptive zones occupied by each morph.

## Conclusions

Color polymorphisms are widespread in nature, with many ecological processes acting to maintain morphs within populations (2). A body of theory and empirical evidence suggests that polymorphism can be both a source of standing phenotypic variation as well as an evolutionary constraint on optimization and diversification (7, 8, 58). However, few studies have brought together morphometric data and well-resolved phylogenies to estimate rates of morphological evolution and speciation to directly test for evolutionary constraints, or lack thereof, in polymorphic lineages versus fixed monomorphic lineages. *Plethodon* are well known for their dorsal stripe polymorphism, and we show that phenotypic optima indeed differed between species of either morph and that unstriped species met the expectations of faster rates of phenotypic evolution. However, our results stand in contrast to other studies that have demonstrated faster speciation rates in polymorphic species (5, 9, 10). In *Plethodon*, we found polymorphic species have relatively slower speciation rates compared to unstriped species or large-bodied species lacking the polymorphism. Our results suggest a strong context-dependence to both rates of phenotypic evolution and species diversification following the fixation of a morph, with asymmetries that can result from alternative adaptations of each morph. Phenotypic evolution may also occur along other axes that we did not consider here, such as physiological tolerance and habitat specialization (59, 60). Investigation of these axes of divergence may shed light on the striped/unstriped polymorphism, as well as diversification within related cryptic species complexes that have since lost the polymorphism. Further experimental work within species, as well as comparative work to establish the shared or contrasting causes of retained polymorphism, will provide valuable insight into processes maintaining polymorphism across the tree of life.

## Materials and methods

### Taxon sampling

There are 57 extant species of *Plethodon*, plus one possible extinct species described from a single specimen (*P. ainsworthi*) (61, 62). *Plethodon* have traditionally been separated into species groups (i.e. hypothesized clades) (Fig. 1; SI Appendix, Table S1) (63). We sampled all extant species, including at least two individuals for most species, except that in western *Plethodon* seven species were each represented by a single sample. Our dataset included 194 samples of *Plethodon* (1–11 samples/species; details of taxon sampling in SI Appendix). We included four outgroup samples, all of which are members of the subfamily Plethodontinae: *Aneides ferreus, Desmognathus apalachicolae, Ensatina eschscholtzii platensis*, and *E. e. xanthoptica*.

We sampled all members of the *P. wehrlei* species complex, representing the most recently described species of *Plethodon* (33, 35, 39). For phylogenetic analyses in which we assigned samples to species, we recognized phylogeographic diversity within some species as separate “species” (SI Appendix, Table S1), which were later reduced to a single sample per recognized species for comparative analyses (see below). In *P. cinereus* and *P. kentucki*, phylogroups included 1–3 samples each (53, 64). Finally, we recognized allopatric phylogroups within two wide ranging species: *P. serratus* (65, 66) and *P. websteri* (67). Further phylogeographic diversity exists within other species (68–70), but we focused on groups for which we had sufficient intraspecific sampling and were supported in preliminary phylogenetic analyses.

### Anchored hybrid enrichment

Details of genomic data collection are provided in the SI Appendix. Briefly, samples were processed using an anchored hybrid enrichment (AHE) protocol (71–73). After processing, we recovered 282 loci with a total of 397,845 bp (mean 1,411 bp per locus; SI Appendix, Table S2). Overall missing data was low (< 30% missing data for 94% of species), and high missing data within samples was almost exclusively in either western *Plethodon* or outgroup samples (SI Appendix, Table S1).

### Phylogenomic analyses and time calibration

As a first pass of the phylogenetic relationships among samples, we estimated a tree of concatenated data using IQ-Tree 2.2.2.6 (74). The model of evolution for each locus was selected using the built-in program ModelFinder (75) with each locus treated as a partition, and node support was inferred using 1,000 ultrafast bootstrap replicates. We also used IQ-Tree in the same manner to estimate a gene tree for each locus.

Gene trees were then used to estimate a species tree with wASTRAL (76), a quartet-based summary method which weighs the contribution of gene trees by branch lengths and support values. For wASTRAL, all 198 samples were assigned to their respective species or phylogroups. For species that were found to be polyphyletic in the concatenated IQ-Tree analysis (*P. shermani, P. glutinosus, P. mississippi, P. metcalfi*), each sample or geographic cluster of samples was treated as an individual tip (SI Appendix, Table S1). Other species that were rendered paraphyletic in the initial concatenated analysis (*P. sequoyah, P. chattahoochee, P. teyahalee*) were maintained as single species, as paraphyly in the concatenated tree could result from the speciation process (35, 77, 78). As an exception, the paraphyletic samples of *P. albagula* were split into two operational units because they represented distant geographic isolates (>1,000 km; SI Appendix, Table S1) (79). After phylogroup assignments, the analysis included 75 tips within *Plethodon*. We measured branch support with local posterior probabilities and considered values >0.95 to be strongly supported, while values between 0.7 and 0.95 were interpreted as modest support (80).

We used MCMCTree in the program PAML 4.10.6 (81) to estimate a time-calibrated phylogeny (details in SI Appendix). Briefly, we fixed the wASTRAL topology, and for each species or phylogroup (including separate tips from polyphyletic species, as described above), we selected the individual with the least missing data as a representative sample. After estimating the substitution rate and gradient and Hessian of the likelihood surface, we estimated divergence dates. As only a single fossil date was available for *Plethodon* (>25 Mya for the root) (82, 83), we used a secondary calibration of the root based on mean estimates across four methods in a previous study of Plethodontidae (37.1–45.3 Mya) (84), setting the 95% prior interval of a skew-normal distribution between these values.

For downstream comparative analyses, we pruned phylogroups, and for species that had paraphyletic or polyphyletic samples, we retained the sample that we considered the best representative of that species based on geography and topology in our estimated phylogenies (described in SI Appendix). We note that these latter cases occurred only in clades of the *glutinosus* group that did not vary in dorsal color morph, and thus sample selection will have little effect on our comparative methods.

### Evolution of color polymorphism

For analyses of the dorsal stripe polymorphism, we used discrete species classifications of striped (S), unstriped (U), or polymorphic (S+U). Most large-bodied species in the *glutinous* group and *wehrlei* species complex (which are not striped/unstriped) were considered a separate color state (“other”). Some species had uncertainty in their color morph assignment (see SI Appendix; Table S3). Briefly, species in the *glutinosus* group that have a variant of red dorsal coloration were either assigned to the striped or polymorphic state (Fig. 1), or they were considered “other” along with the remaining *glutinosus* group. Three species in western *Plethodon* have an ontogenetic shift in color morph that commonly results in adults losing the dorsal stripe. We tested *a priori* assignments for these species (Fig. 1), as well as schemes where each was considered striped, unstriped, or polymorphic. These combinations resulted in eight total classification schemes.

We obtained species-level morphometric data for 52 species from Baken and Adams (2019) and 37 species from Adams et al. (2009). A weighted average of each measurement across the two studies was used for species present in both datasets. Data for missing species were compiled from additional sources (see SI Appendix). Morphometric measurements included snout-vent length (SVL); head length (HL); snout-eye distance (SE); forelimb length (FLL); and hindlimb length (HLL). Tail length (TL) was not included here because tail breakage and regrowth can cause high variation among specimens. To account for body size variation, we divided HL, SE, FLL, and HLL by SVL and log-transformed the values, focusing on changes in relative proportions (85, 86). We also included median costal groove counts (CG) from Fisher-Reid and Wiens (2015) and other sources (34, 87–90) as a metric of relative trunk elongation, as the number of CG is one fewer than the number of trunk vertebrae (91). We measured phylogenetic signal in each morphometric trait using Pagel’s lambda (92) and Blomberg’s K (93) and we estimated a phylogenetic PCA (pPCA) of morphometric characters (94) using the R package ‘phytools’ (95).

We tested 12 models of discrete character evolution, 7 of which included an explicit polymorphism parameterization in which transitions between the striped and unstriped states needed to pass through the polymorphic state (symmetrically or asymmetrically; SI Appendix, Fig. S3). For the fourth character state for the eastern large-bodied *Plethodon* (“other”), we assumed that transitions to “other” occurred from the polymorphic state at a separate rate, and “other” could transition to either S or S+U at another rate to account for striped or polymorphic species nested within clades classified as “other.” To estimate ancestral states, we performed stochastic character mapping, incorporating model uncertainty by simulating all models in proportion to their AIC weights for a total of 1,000 simulations. To accommodate uncertainty in color morph classifications, we repeated this procedure for the eight color classification schemes that varied in combinations of how some western *Plethodon* were assigned and whether some members of the *glutinosus* group were assigned as “other” or to one of the three color morph states.

Finally, we tested whether different color morph states had different evolutionary rates or optima using the R package ‘OUwie’ (96). We compared the fit of a single rate Brownian Motion model (BM1), a multi-rate BM model (BMS), a single optimum Ornstein–Uhlenbeck model (OU1), a multi-optima OU model (OUM), and a multi-rate and optima OU model (OUMV). A single sample of 100 stochastic character maps from each color classification scheme was used as ancestral states in separate analyses using the BMS, OUM, and OUMV models, whereas BM1 and OU1 are not affected by trait state. Morphometric response variables mostly captured relative elongation. We used the first phylogenetic PCA axis (pPC1) as a metric of elongation, but we also tested univariate metrics of total size-corrected limb length, size-corrected head length, and raw count of costal grooves to ensure results were not biased by principal components (97). The distribution of AIC scores across stochastic character maps was used to evaluate support for different models. We report mean rates of evolution (σ^2^) and trait optima (θ) across OUMV models (see Results) along with standard errors given the variance across stochastic character maps and color classification schemes.

### Diversification rates

We estimated lineage-specific diversification rates, or tip rates, using the DR statistic, λ_DR_ (98), which we calculated using the R package ‘epm’ (99). We used λ_DR_ as a simple metric of diversification instead of more sophisticated state-based models (e.g. MuHiSSE; 100) due to the relatively small size of our phylogeny. λ_DR_ tends to be stronger predictor of speciation rates than net diversification rates (101); therefore, we interpret results as corresponding to speciation rates. We calculated mean λ_DR_ among species within color morph states for each color classification scheme.

## Supporting information

Phylogenetic Trees

## Data availability

Phylogenies are included as Supplemental Data. All data associated with this project will be made available upon publication.

## Acknowledgements

We are indebted to Richard Highton for the use of many genetic samples, and whose extensive work with *Plethodon* made the phylogenomic tree possible. We also thank Bryan Carstens for contributing a sample of *P. idahoensis*, and Jessica Oswald for help with molecular work. Tara Pelletier, Carl Anthony, and Brendan Enochs provided helpful input on color morph classifications, and Zach Felix and Dean Adams generously shared morphometric data. We thank Michelle Kortyna at Florida State University’s Center for Anchored Phylogenomics for assistance with data collection. This work was supported by an NSF Postdoctoral Research Fellowship in Biology to M.M. Hantak (NSF DEB 2010776), the University of Florida to D.C. Blackburn, and Ohio University to S.R. Kuchta.

