## Supplementary Appendix for "Loss of a morph is associated with asymmetric character release in a radiation of woodland salamanders"

**This PDF file includes:**

Supporting text
- Taxon sampling
- Anchored hybrid enrichment
- Phylogenomic analyses (Figure S1 – S2)
  - Timetree estimation
- Color classification schemes
- Ancestral character reconstruction (Figure S3 – S4)
- Morphometrics
- Variation in phylogenetic PCA (Figure S5)
- Rates of evolution and evolutionary optima (Figure S6 – S7)

SI References

**Other supporting materials for this manuscript include the following:**

Table S1 – Sample information
Table S2 – Per-locus summary statistics
Table S3 – Morphometric raw data and discrete color classifications
Table S4 – Discrete character evolution model rankings
Table S5 – Morphometric phylogenetic signal
Table S6 – Phylogenetic PCA (pPCA) loadings

Tree files:
- “plethodon-iqtree.tree” – IQ-Tree treefile
- “plethodon-wastral.tree” – wASTRAL treefile
- “plethodon-timetree-unpruned.tree” – MCMCTree timetree (unpruned)
- “plethodon-timetree-pruned.tree” – MCMCTree timetree (pruned)

### Supporting Information Text

#### Taxon sampling

*Plethodon* consists of 57 extant species across four major clades: 1) western *Plethodon* (N = 9 species; also assigned to the subgenus *Hightonia*); 2) the *cinereus* group (for clarity, we omit the genus name when referring to species groups throughout) (N = 10); 3) the *wehrlei-websteri* group, also sometimes referred to as the "*wehrlei-welleri* group" (1) (N = 10); and 4) the *glutinosus* group (N = 28). As the names suggest, western *Plethodon* are found in western North America, whereas all other groups are found in eastern North America, comprising "eastern *Plethodon*" (subgenus *Plethodon*). We included samples from all extant species. Phylogenetic inference has been most challenging in eastern *Plethodon* (1–4), and we made an effort to sample multiple individuals per species across different localities (Supplementary Table S1).

Within two species, we recognized phylogroups identified in recent work with the same samples and genetic data. *Plethodon cinereus* included five groups ("Groups 1 – 5"), which were subsampled here using 1 – 3 samples per group (5). *Plethodon kentucki* included four groups ("Cumberland," "Kentucky," "Kanawha," "New"), subsampled here using 3 samples per group (6). For some other species, all samples were retained and separated into geographic clusters: *Plethodon serratus* ("Appalachia," "Missouri," "Louisiana," "Ouachita") (7, 8); *Plethodon websteri* ("AL," "MS," "GA," "SC") (9); *Plethodon albagula* ("Texas," "Ozark") (10). Some other species were not separated into multiple geographic groups *a priori*, but instead multiple geographic groups were recognized after a concatenated analysis of individuals in IQ-Tree (see below and main text): *Plethodon metcalfi* ("N," "S"); *Plethodon shermani* ("E," "W"); *Plethodon glutinosus* ("N," "S"); *Plethodon mississippi* ("E," "W"). We recognize all recently described or resurrected species of the *P. wehrlei* species complex: *Plethodon dixi*, *P. jacksoni*, and *P. pauleyi* (11–13).

#### Anchored hybrid enrichment

Genomic DNA was extracted from blood or tail tissue using Qiagen DNeasy blood and tissue kits (Qiagen Corp.). Samples were processed at the Florida State University Center for Anchored Phylogenomics using an anchored hybrid enrichment (AHE) protocol (14). Each sample was standardized to ~20 ng/μL and sonicated to an average fragment size of 150 – 300 bp using a Covaris E220 focused ultrasonicator. Library preparation was performed using a Biomex FXp liquid-handling robot (Beckman Coulter) following (15). Samples were indexed and enriched using an Agilent Custom SureSelect Kit with amphibian-specific probes (16). Samples were then pooled for sequencing on a PE150 lane of a HiSeq 2000 DNA sequencer (Illumina). Data were processed using scripts and methodology following (17–19), except where noted. Briefly, overlapping reads were merged and assembled. We did not phase alleles for this study. Low-level contaminants were filtered, and we performed an initial assessment of orthology based on pairwise distances for each locus. We aligned sequences using MAFFT (20), which were trimmed and masked to remove sites with excessive missing data (18). Within 20 bp windows, all sites were masked if they contained fewer than 14 "good" sites (i.e. where the most common base character was present in >50% of samples). Sites with masked or missing bases in >50% of samples were removed (18). We retained 282 AHE loci for this study. Sequence statistics per locus can be found in Supplementary Table S2, and the proportion of total missing data per sample is found in Supplementary Table S1.

### Phylogenomic analyses

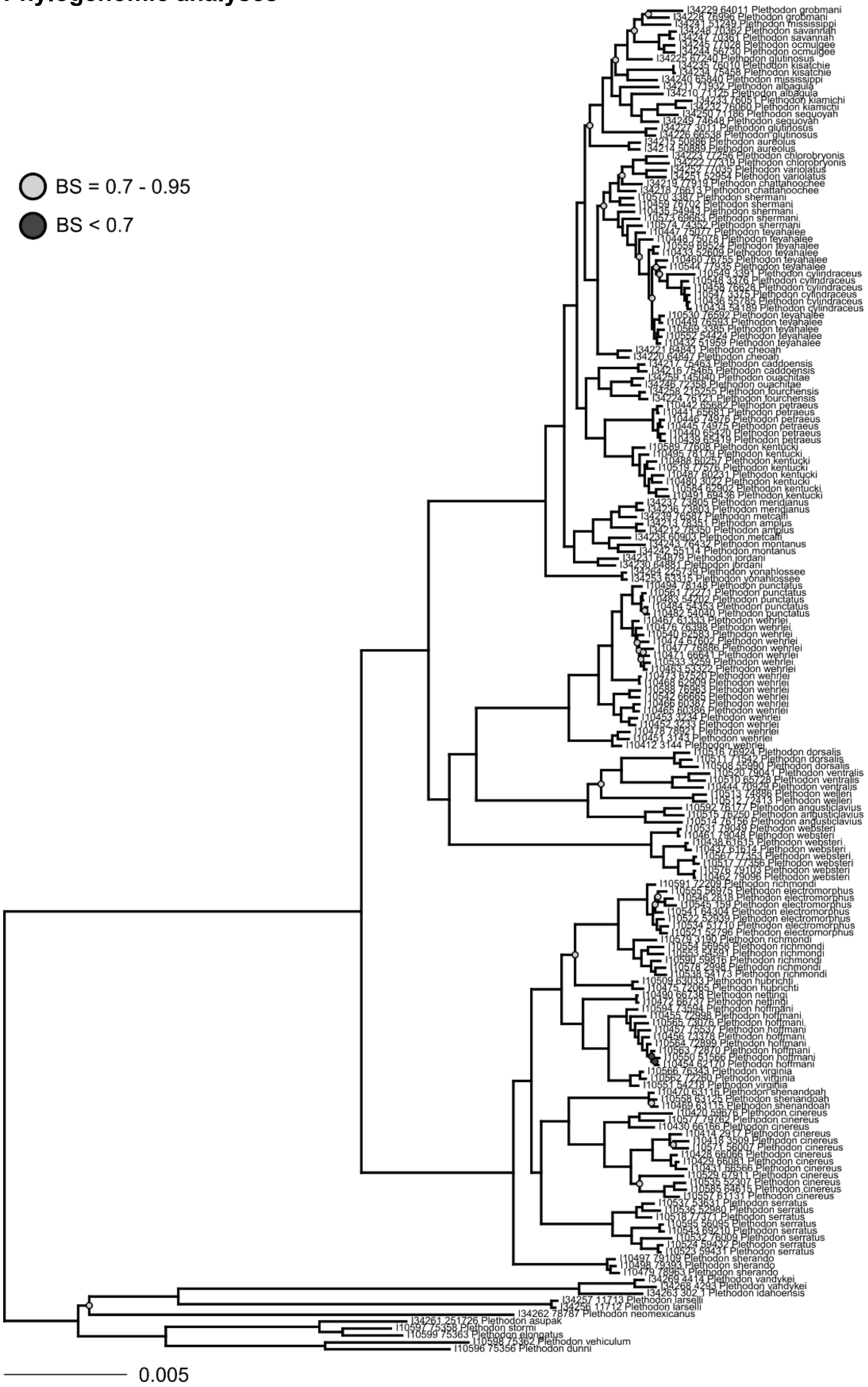


**Supplementary Figure S1**. Concatenated phylogeny of all samples of *Plethodon* estimated using IQ-Tree. Outgroups have been pruned for visualization. Node support represents bootstrap support, with unlabeled nodes having values >0.95. Sample names can be matched with updated species assignments (e.g. within the *P. wehrlei* species complex) and phylogroup assignments used for species tree estimation in Supplementary Table S1.

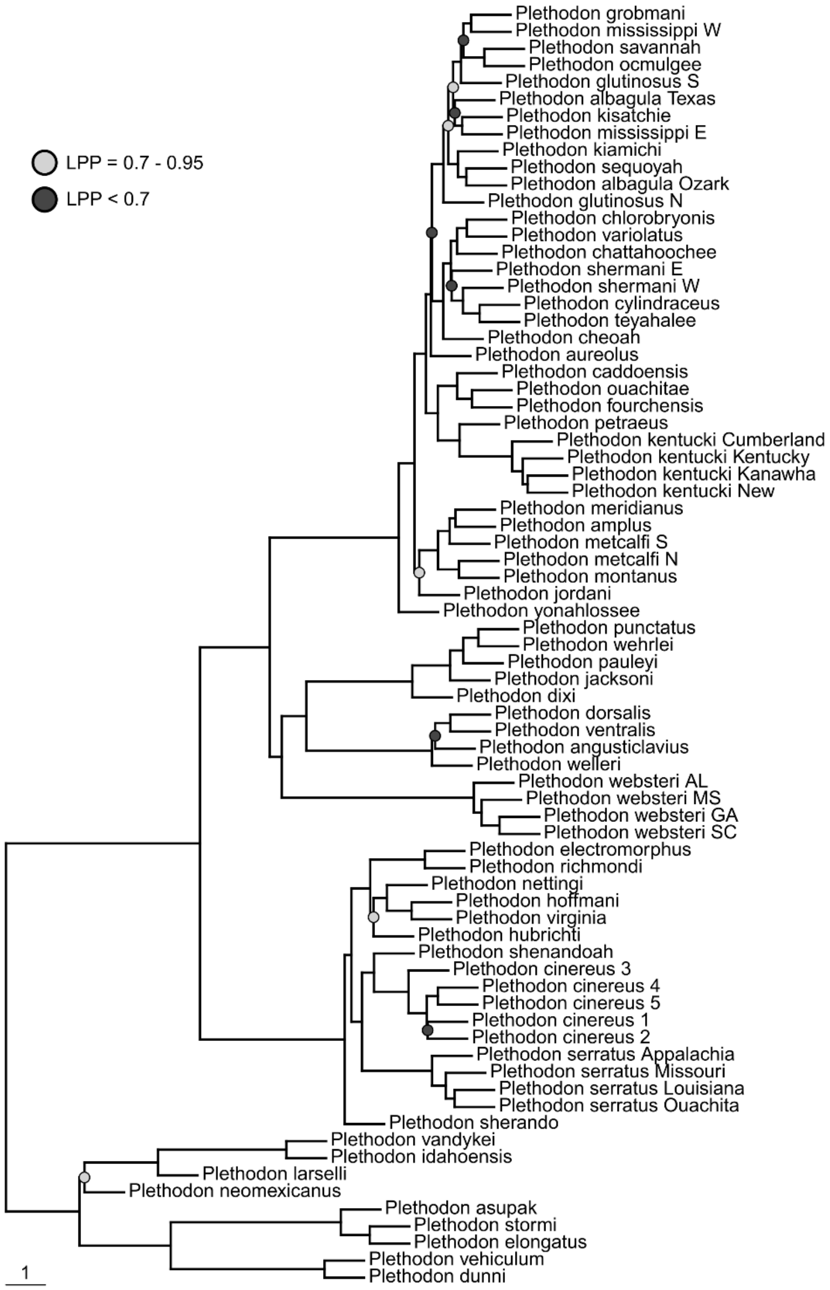


**Supplementary Figure S2**. Species tree of *Plethodon* estimated using wASTRAL. Outgroups have been pruned for visualization. Node support represents local posterior probabilities, with unlabeled nodes having values >0.95. Branch lengths represent coalescent units, with external branches set to 1. Samples were assigned to species or phylogroups based on concatenated analysis of individuals (Supplementary Table S1).

**Timetree estimation.** We estimated a timetree using the program PAML 4.10.6 (21). We initially attempted Bayesian analysis with starBEAST3 (22), using a partitioned analysis of the 20 longest loci with minimal missing data (fewer than 10% of samples with completely missing data) that were more “clocklike” (coefficient of variation in root-to-tip distance less than the median value across loci) (23); however, this analysis had ESS <100 for several parameters (typically population sizes, tree heights, and tree distances) after a total of 400,000 iterations across two independent chains, suggesting the analysis would require fewer loci, simpler models, or much longer runs to converge. Therefore, we estimated divergence times using MCMCTree. For this analysis, the species tree topology inferred using wASTRAL was fixed, and for each species or phylogroup—including separate tips from polyphyletic species—we selected the individual with the least missing data as a representative sample. We first obtained a rough estimate of the overall substitution rate across loci using BASEML, treating loci as partitions. The root node was fixed at 40 Mya, a date estimated for the split between eastern and western *Plethodon* in previous analyses (Shen et al. 2016; see below), and we assumed a GTR+ Γ model for each locus. The estimated substitution rate was 5.335 x $10^{-3}$ substitutions per 10 Ma, which was subsequently used to parameterize a broad gamma-Dirichlet distribution of locus rates (rgene_gamma = 1 382.40921). Next, we estimated the branch lengths as well as the gradient and Hessian of likelihood surface (usedata = 3). Finally, we used the estimated branch lengths and a fossil calibration of the root node to obtain divergence dates (usedata = 2). Few fossils are available within *Plethodon*, but we were able to constrain the root to be at least 25 Mya based a late Oligocene fossil from Montana consistent with western *Plethodon* (25–27). Including this same fossil calibration, Shen et al. (2016) used 50 loci to estimate divergence dates for Plethodontidae, with mean divergences between eastern and western *Plethodon* ranging from 37.1 – 45.3 Mya across four analyses. We used these values to set a skew-normal distribution on the root of *Plethodon* which included these values in the 95% the prior probability [SN(4.0, 0.2377331, 0.7183491)]. We used a birth-death model with a sampling proportion of 0.9 (i.e. assuming most species have been sampled). MCMC chains were set to collect samples every 100 iterations for a total of 1,000 samples, following a burn-in of 10,000 iterations. Chains were evaluated using Tracer V1.7.2 (28) to ensure convergence and ESS >200.

For downstream comparative analyses that required species to be represented by a single tip, polyphyletic or paraphyletic species were reduced to the unit that we inferred to be the best representative of that species based on geography and topology in our estimated phylogenies: 1) the Texas isolate of *P. albagula*; 2) the northern (N) group of *P. glutinosus*; 3) the southern (S) sample of *P. metcalfi*; 4) the eastern (E) sample of *P. mississippi*; and 5) the eastern (E) sample of *P. shermani* from the Wayah isolate. In general, we prioritized groups that were less likely to occur at a contact zone with other species.

**Color morph classification schemes**

Species were classified into color morph categories based on species accounts (29, 30) and supplemented with expert opinion (see acknowledgements) and photographs from AmphibiaWeb (30) or iNaturalist (31). Many species had straightforward color morph classifications, but two areas of the phylogeny presented challenges that we addressed by exploring multiple classifications. First, eastern large-bodied *Plethodon* (the *glutinosus* group and the *wehrlei* species complex) generally lack a stripe, yet their patterning is distinct from other unstriped species. *Plethodon yonahlossee*, *P. petraeus*, and *P. ouachitae* are exceptions, each with a red dorsum (a possibly polymorphic feature for *P. ouachitae*), but their stripe is conspicuously different in width, pattern, and hue than any other species, and may have a different developmental basis or functional role. We tested classifications in which 1) most members of the *glutinosus* group and *wehrlei* complex were assigned to a fourth category consistent with eastern large-bodied *Plethodon* ("other") with the exception of the three striped or polymorphic species listed above; or 2) all members of these clades were all classified as "other". Second, western *Plethodon* classifications were complicated by an ontogenetic change observed in *P. neomexicanus*, *P. asupak*, *P. stormi*, and *P. elongatus*, in which some or all individuals are striped as juveniles, but the stripe gradually dulls in adulthood to be indistinguishable from an unstriped morph. The degree to which the stripe dulls varies among individuals and species. In previous work, *P. neomexicanus* was considered unstriped, while *P. elongatus* was considered striped, and the remaining two species were not sampled (32). Ontogenetic change is usually not considered polymorphism, and species are typically classified based on their adult morphology (33); however, because there is little data regarding individual/population variation and the duration of the stripe into adulthood, we tested four alternative classifications based on photographs available on iNaturalist (31) and AmphibiaWeb (30) and correspondence with other scientists (see Acknowledgements): *Plethodon neomexicanus* was always considered unstriped, while 1) *P. asupak* and *P. stormi* were considered unstriped, and *P. elongatus* striped; or these three species were all considered 2) unstriped; 3) striped; or 4) polymorphic. In total, we tested 8 classification schemes. We were interested in the way in which these different classifications affected ancestral state reconstructions and estimates of evolutionary rates and optima.

Most of our color classifications matched those of Fisher-Reid and Wiens (2015). However, we considered *P. hubrichti* unstriped because its golden stripe better resembles a dense concentration of iridophores rather than a red stripe. We also assigned *P. dunni* as polymorphic due to the presence of unstriped individuals and populations, once considered a separate species (*P. gordoni*) (34). We categorized *P. petraeus* as either striped or “other," depending on the classification scheme. Finally, species not included in Fisher-Reid and Wiens (2015) included *P. sherando* (polymorphic), newly described species of the “*P. wehrlei* complex” (all “other”), and *P. asupak* and *P. stormi* (varied across classification schemes).

**Ancestral character reconstruction**

We tested 12 models of discrete character evolution for each color morph classification scheme. These models tested multiple hypotheses regarding the directionality of color morph evolution, while also accounting for transitions to and from the large-bodied color states (“other”), which were held constant across models (Supplementary Fig. S3). Models were fit using ‘fitMk’ in ‘phytools’ (35). Although ancestral character reconstructions for eastern *Plethodon* were generally consistent across schemes, the root state and the states within western *Plethodon* were highly variable (Supplementary Fig. S4), suggesting sensitivity to the color classifications within these groups.

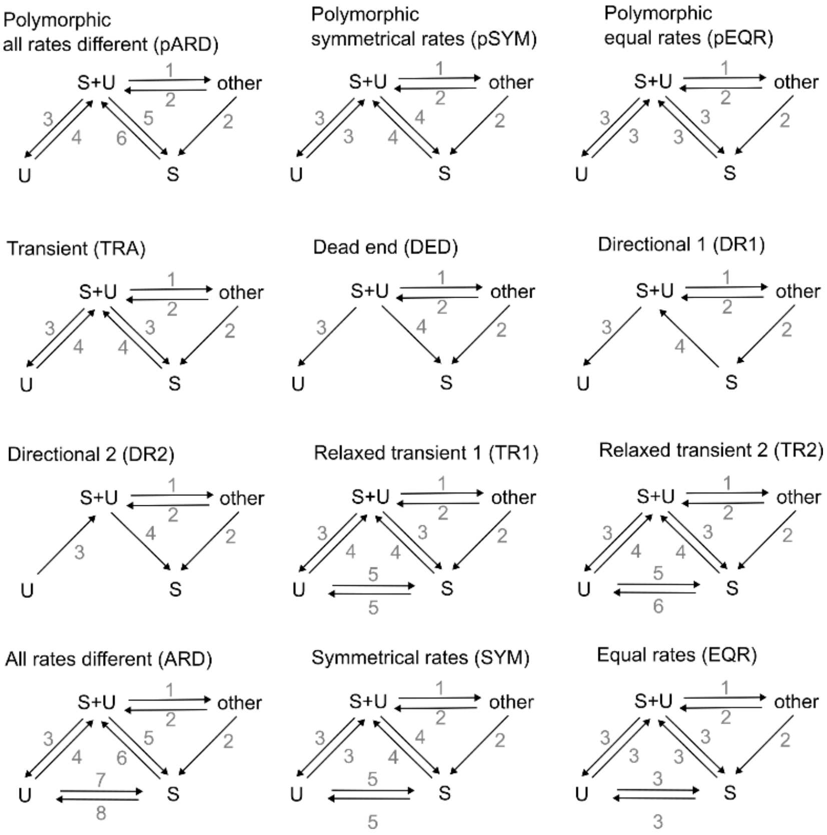


**Supplementary Figure S3**. Models of discrete character evolution for color polymorphism (S = striped, U = unstriped, S+U = polymorphic). Arrows indicate which parameters were included in the model, and arrows with the same number indicate shared parameter values.

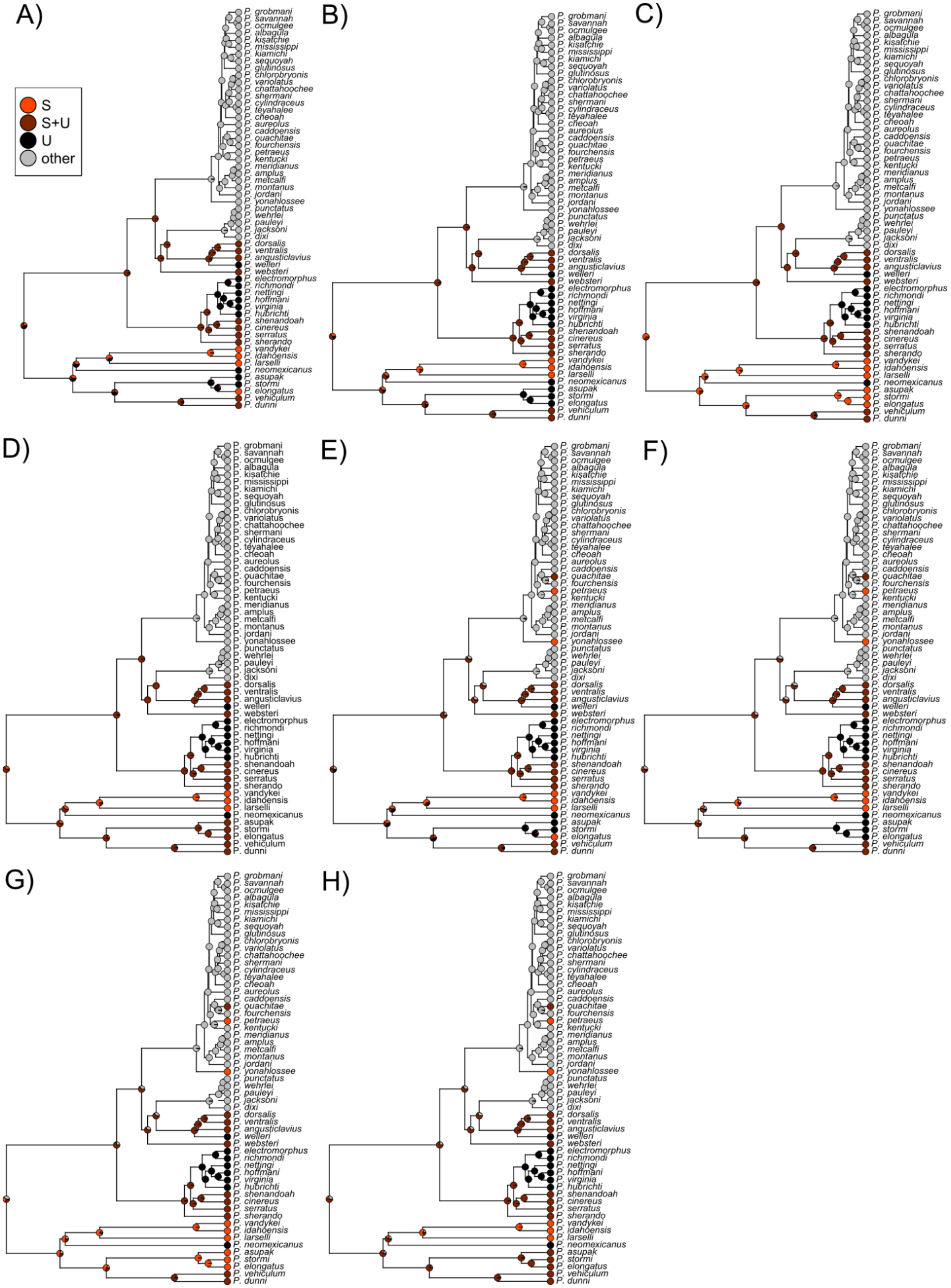

A)
B)
C)
D)
E)
F)
G)
H)
S
S+U
U
other
P. grobmani
P. savannah
P. ocmulgee
P. albagula
P. kisatchie
P. mississippi
P. kiamichi
P. sequoyah
P. glutinosus
P. chlorobryonis
P. variolatus
P. chattahoochee
P. shermani
P. cylindraceus
P. teyahalee
P. cheoah
P. aureolus
P. caddoensis
P. ouachitae
P. fourchensis
P. petraeus
P. kentucki
P. meridianus
P. amplus
P. metcalfi
P. montanus
P. jordani
P. yonahlossee
P. punctatus
P. wehrlei
P. pauleyi
P. jacksoni
P. dixi
P. dorsalis
P. ventralis
P. angusticlavius
P. welleri
P. websteri
P. electromorphus
P. richmondi
P. nettingi
P. hoffmani
P. virginia
P. hubrichti
P. shenandoah
P. cinereus
P. serratus
P. sherando
P. vandykei
P. idahoensis
P. larselli
P. neomexicanus
P. asupak
P. stormi
P. elongatus
P. vehiculum
P. dunni

**Supplementary Figure S4.** Ancestral character reconstruction of color morph state using stochastic character mapping (SCM) for eight color classification schemes (A – H; scheme E is the same as Figures 1 and 3a in the main text). Schemes A – D considered all members of the *glutinosus* group and *wehrlei* species complex to be a separate state ("other"), while E – H allowed some species to be striped (S) or polymorphic (S+U); schemes further varied in the states of *P. asupak*, *P. stormi*, and *P. elongatus*. For each scheme, 1,000 SCMs were performed across models of discrete character evolution in proportion to their AIC weights.

**Morphometrics**

We obtained morphometric data for most species from Baken and Adams (2019) (N = 52) or Adams et al. (2009) (N = 37). Measurements included snout-vent length (SVL), head length (HL), snout-eye distance (SE), forelimb length (FLL), and hindlimb length (HLL). Tail length (TL) was not included here because tail breakage and regrowth can cause high variation among specimens. A weighted average of each measurement based on sample size was used as the species value. Species without data in these studies were the newly described members of the *P. wehrlei* species complex (11–13) and *P. sherando* (38). For the *P. wehrlei* species complex, we were able to obtain values for *P. wehrlei*, *P. dixi*, and *P. jacksoni* from (37) based on localities of the collected specimens and new species designations (11, 12) (D. Adams, personal communication). Most assignments were straightforward; however, the samples we included in *P. jacksoni* were from a geographic isolate near known *P. jacksoni*, but this isolate was not sampled in genetic studies. For *P. pauleyi,* which had no specimens near known localities, we used mean values of SVL, BW, FLL, and HLL from the holotype and allotypes (N = 4) (11). HL and SE were not used because they were unmeasured or measured differently across studies. Similarly, for *P. sherando* we used midpoint measurements of the holotype and allotype from Highton (2004) for SVL, FLL, and HLL. Costal groove counts were obtained from Fisher-Reid and Wiens (2015). For species without CG data in that study, we used other published values (38–41). We also renumbered *Plethodon sequoyah* as 16 CG based on Table 11 in (42). Because missing morphometric data that remained were limited to a small number of traits in *P. pauleyi* and *P. sherando*, we used the R package 'Rphylopars' for phylogenetic imputation (43).

**Variation in phylogenetic PCA**

Results of the phylogenetic PCA (pPCA) with species labeled are provided in Fig. S5, which also highlights species with alternative color classifications. Color morphs tended to cluster together. One western *Plethodon* species, *P. elongatus*, included uncertainty in color classification, but when scored as striped (the rightmost light-red triangle in Supplementary Fig. S5), it appeared more elongated than other striped species.

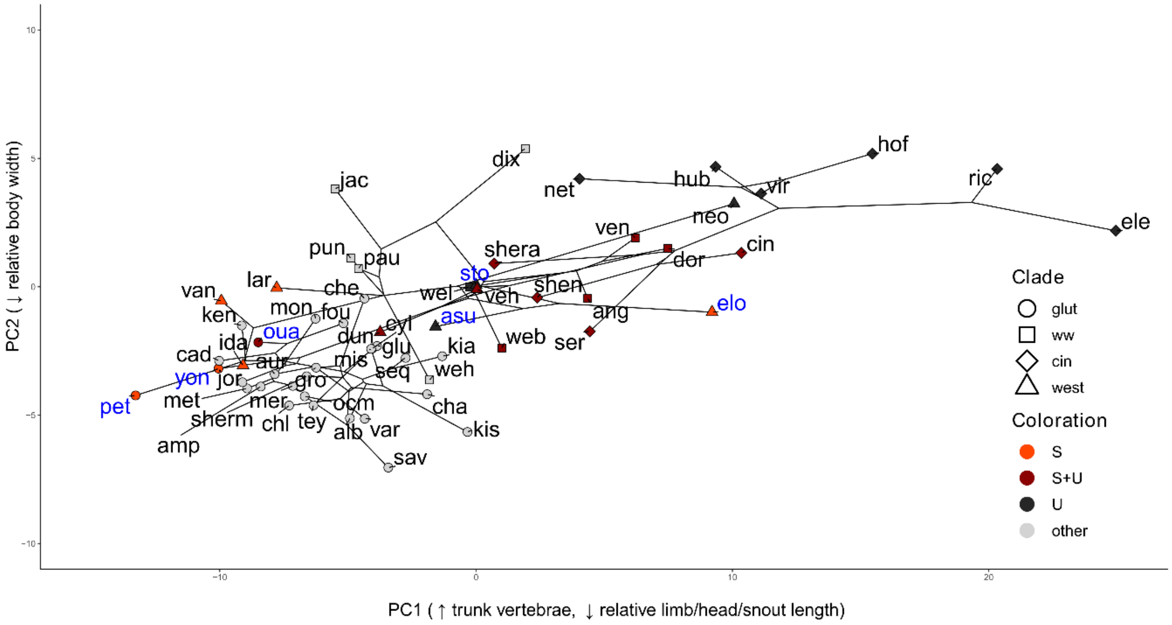


**Supplementary Figure S5.** Phylogenetic PCA of morphometric data. Results match Fig. 3a of the main text, but species are identified with the first 3–5 letters of their specific epithet. Blue species labels represent species with alternative color classifications. The first axis is consistent with increasing numbers of trunk vertebrae and decreasing body-size-corrected lengths of the limbs, head, and snout. The second axis represents decreasing size-corrected body width. Points are labeled by species groups and colored by dorsal color state (S = striped, S+U = polymorphic, U = unstriped). Represented species color classifications match Fig. 1.

### Rates of evolution and evolutionary optima

We focused on OUMV models, which generally had better AIC scores (Supplementary Fig. S6) and included both parameters of interest: the rate of evolution ($\sigma^2$) and the evolutionary optimum ($\theta$). Our interpretation of greater relative body elongation in the unstriped morph and lower in the striped morph, and faster rates of evolution in the unstriped morph are supported by analyses using the univariate metrics of CG count, total size-corrected limb length (i.e. the sum of log-transformed, SVL-corrected FLL and HLL), and size-corrected HL (Supplementary Fig. S7).

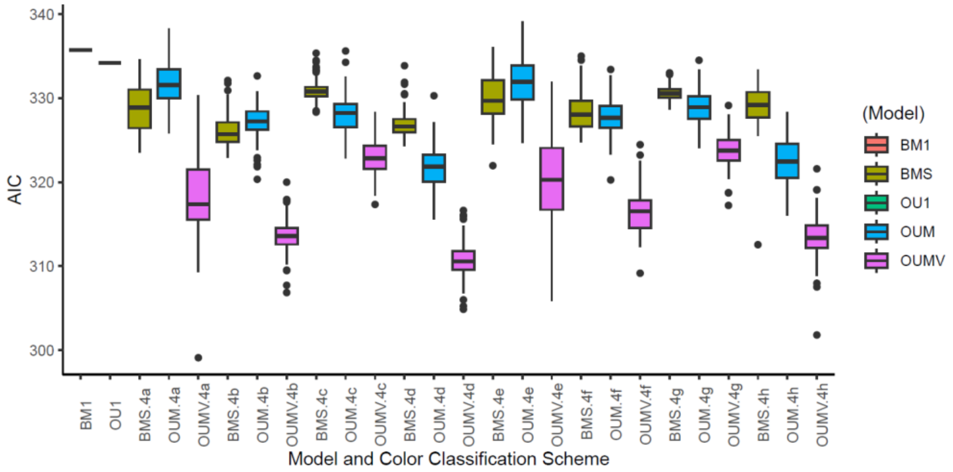

**Supplementary Figure S6**. Distribution of OUwie AIC values across color classification schemes and evolutionary models for pPC1 (variation in numbers of trunk vertebrae and body-size-corrected lengths of the limbs, head, and snout), with each combination represented by 100 stochastic character maps. BM1 and OU1 (leftmost boxplots) did not depend on color morph state, while the remaining models allowed the four color morph states (4a–4h) to vary in either evolutionary rate (BMS), optima (OUM), or both (OUMV).

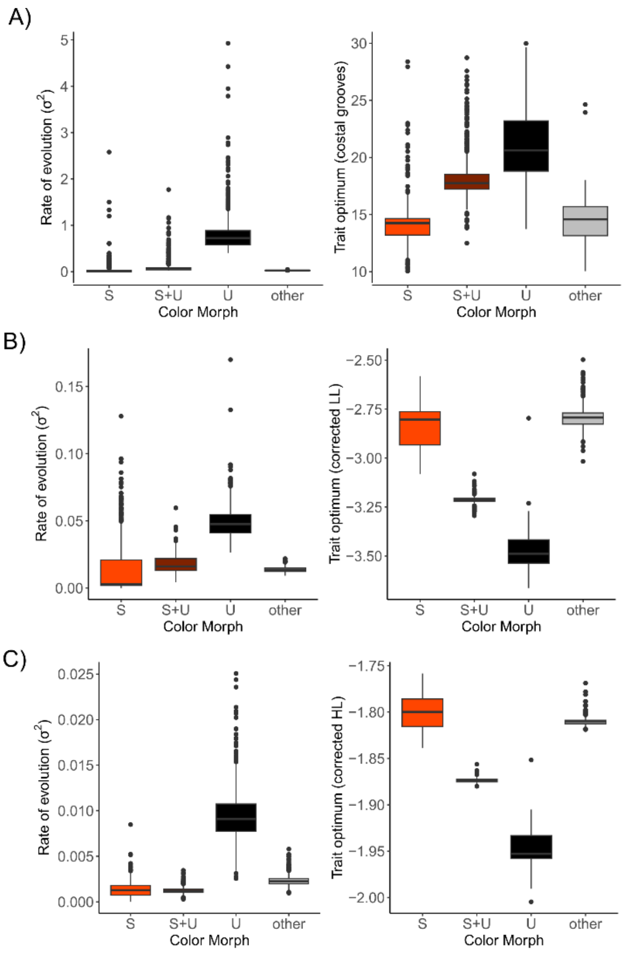


**Supplementary Figure S7**. OUwie results, including evolutionary rates (*left*) and optima (*right*), for alternative metrics of body elongation: A) Costal groove count, B) total log-transformed, size-corrected limb length (i.e. hindlimb plus forelimb), and C) log-transformed, size-corrected head length. Results for each color morph state represent pooled values across color classification schemes of multi-rate and optima OU models (OUMV), given 100 stochastic character mapping per scheme and model. For costal groove count (A), 362 outliers (out of 3,200 total values) with unrealistically high or low estimated optima (>30 or <10) were excluded and are not visible.
